# Environmental DNA enables detection of terrestrial mammals from forest pond water

**DOI:** 10.1101/068551

**Authors:** Masayuki Ushio, Hisato Fukuda, Toshiki Inoue, Kobayashi Makoto, Osamu Kishida, Keiichi Sato, Koichi Murata, Masato Nikaido, Tetsuya Sado, Yukuto Sato, Masamichi Takeshita, Wataru Iwasaki, Hiroki Yamanaka, Michio Kondoh, Masaki Miya

## Abstract

Terrestrial animals must have frequent contact with water to maintain their lives, implying that environmental DNA (eDNA) originating from terrestrial animals should be detectable from places containing water in terrestrial ecosystems. Aiming to detect the presence of terrestrial mammals using forest water samples, we applied a set of universal PCR primers (MiMammal, a modified version of fish universal primers) for metabarcoding mammalian eDNA. After verifying the primers’ usefulness *in silico* and using water samples from zoo cages of animals with known species compositions, we collected five 500-ml water samples from ponds in two cool-temperate forests in Hokkaido, northern Japan. Using eDNA extracted from the water samples, we constructed amplicon libraries using MiMammal primers for Illumina MiSeq sequencing. MiMammal metabarcoding yielded a total of 75,214 reads, which we then subjected to data pre-processing and taxonomic assignment. We thereby detected species of mammals common to the sampling areas, including deer (*Cervus nippon*), mouse (*Mus musculus*), vole (*Myodes rufocanus*), raccoon (*Procyon lotor*), rat (*Rattus norvegicus*) and shrew (*Sorex unguiculatus*). Previous applications of the eDNA metabarcoding approach have mostly been limited to aquatic/semiaquatic systems, but the results presented here show that the approach is also promising even in forest mammal biodiversity surveys.

## I. INTRODUCTION

Environmental DNA (eDNA) is a genetic material that is found in an environment and derived from organisms living in that habitat, and researchers have recently been using eDNA to detect the presence of macro-organisms, particularly those living in an aquatic environment. In the case of macro-organisms, eDNA originates from various sources such as metabolic waste, damaged tissue, or sloughed skin cells [1], and the eDNA contains information about the species identity of organisms that produced it. Ficetola et al. [2] first demonstrated the usefulness of eDNA for detecting the presence of an aquatic vertebrate (American bullfrog invasive in France) in natural wetlands. Subsequently, eDNA in aquatic ecosystems has been repeatedly used as a monitoring tool for the distributions of fish species in ponds, rivers and seawater [3–5] as well as the distributions of other aquatic/semiaquatic non-fish vertebrates such as giant salamanders [6]. Those studies have shown that the use of eDNA can be a promising approach as an efficient non-invasive monitoring tool for biodiversity in aquatic ecology.

Although earlier studies used quantitative PCR and species-specific primers to amplify a particular region of eDNA [2–4, 6, 7], researchers have begun to apply massively parallel sequencing technology (e.g., Illumina MiSeq) and universal primer sets to eDNA studies (eDNA metabarcoding approach) [8, 9]. This ap-proach greatly improved the effiiency and sensitivity of eDNA detection. A previous study demonstrated that an eDNA metabarcoding approach using fish-targeting universal primers (MiFish primers) enabled the detection of more than 230 fish species from seawater in a single study [8]. Accordingly, the eDNA metabarcoding approach has become a cost- and labor-effective approach for capturing aquatic biodiversity.

The use of eDNA is, however, not necessarily limited to aquatic/semiaquatic vertebrates, because terrestrial animals also have frequent opportunities to contact aquatic systems. For example, Rodgers and Mock [7] tested the potential of drinking water for mammals as a medium that contains eDNA originating from terrestrial mammals. They hypothesized that when a terrestrial mammal (coyote; *Canis latrans*) drinks from a water source, DNA from its saliva and mouth tissues is shed and can be used as a tool for species identification. In their study, they successfully amplified a specific region of eDNA using coyote-specific primers. That study demonstrated that eDNA is a potentially useful tool even for detecting and monitoring the presence of mammals living in a terrestrial ecosystem if one can collect appropriate media that contain mammalian DNA. Mammals living in a forest ecosystem often contact/utilize water in a pond [10], and thus, a forest pond may provide an opportunity for researchers to efficiently collect mammals’ eDNA.

In the present study, we hypothesized that, when mammals utilize/contact water, they shed their tissues into the water as a source of eDNA, and that this eDNA can be used to detect the presence of mammals living in a forest ecosystem. To improve the sensitivity and efficiency of the eDNA approach, we modified the previously developed fish-targeting universal primers (MiFish primers) [8] by accommodating them to mammal-specific variation, and conducted mammalian eDNA metabarcoding. The versatility of the primers was tested *in silico*, and their accuracy was further tested by analyzing water samples from zoo cages containing animals of known species composition. Finally, we examined the effectiveness of the new primer set using water samples from field sites.

## II. MATERIALS AND METHODS

### A. New primer sets and test of versatility

Mitochondrial DNA (mtDNA) was chosen as the genetic marker because the copy number per cell of mtDNA is greater than that of nuclear DNA, and the detection rate is therefore expected to be higher when using the former. Our primers were designed by modifying previously developed MiFish primers [8], which corresponded to regions in the mitochondrial 12S rRNA gene, and we named our primers MiMammal-U (forward primer = GGG TTG GTA AAT TTC GTG CCA GC; reverse primer = CAT AGT GGG GTA TCT AAT CCC AGT TTG). In addition, we also designed MiMammal-E (forward primer = GGA CTG GTC AAT TTC GTG CCA GC; reverse primer = CAT AGT GAG GTA TCT AAT CTC AGT TTG) and MiMammal-B (forward primer = GGG TTG GTT AAT TTC GTG CCA GC; reverse primer = CAT AGT GGG GTA TCT AAT CCC AGT TTG) primers after preliminary experiments to accommodate sequence variations in the priming sites of elephants and bears, respectively. The versatility of these primers was tested *in silico* and using extracted DNA and eDNA from zoo cages (see Figs. S1-S2 and Tables S1-S3).

### B. Water sampling and DNA extraction

We collected five water samples from four ponds in two cool-temperate forests in Hokkaido, northern Japan (Teshio Experimental Forest, 44^°^54'55” N, 142°1'21” E; Tomakomai Experimental Forest, 42°39'32” N, 141°36'19” E) to examine the use of the primers for metabarcoding mammalian eDNA from natural forest ponds with unknown mammal compositions in an open ecosystem (Fig.1). All sampling and filtering equipment was washed with a bleach solution before use. Approximately 500-ml water samples were collected from forest ponds using polyethylene bottles (Fig.1).

**Figure 1.**
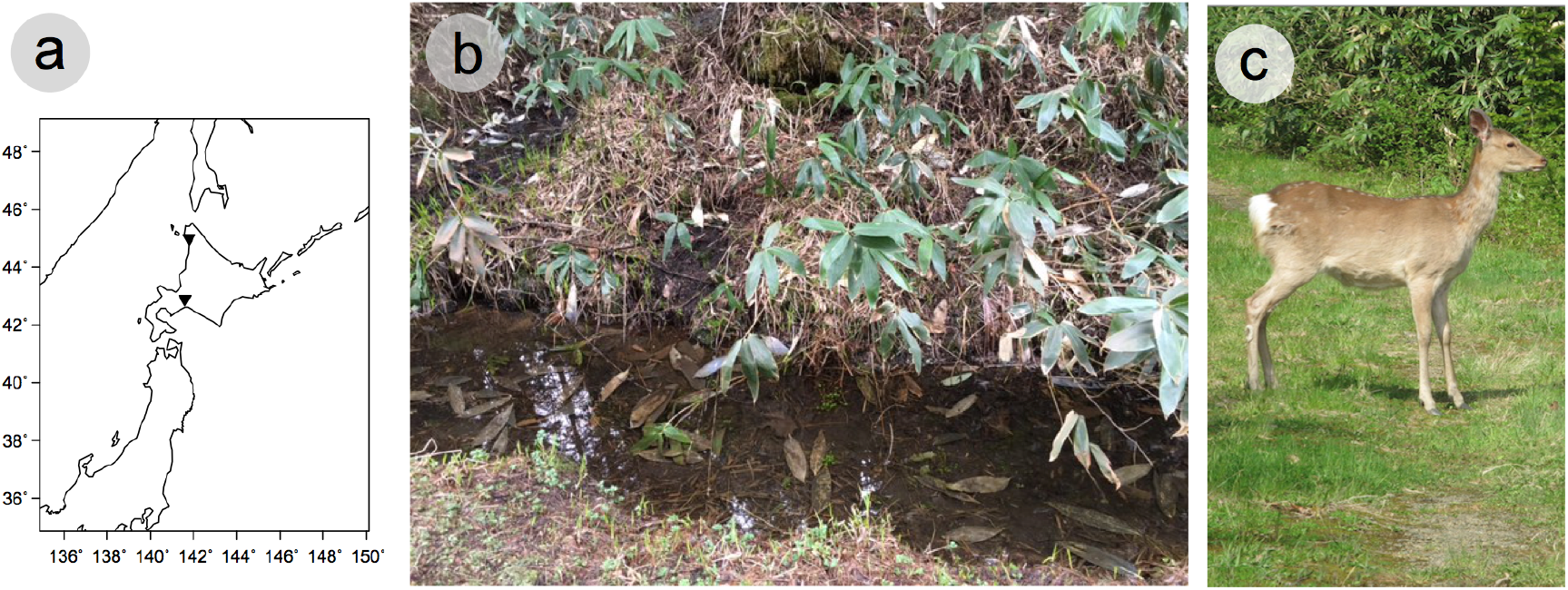
Field survey in natural cool-temperate forests in Hokkaido, Japan. Location of the two cool-temperate forests (a). The upper and lower black triangle indicates Teshio Experimental Forest and Tomakomai Experimental Forest, respectively. One of our sampling ponds in Teshio forest (b). A Sika deer, *Cervus nippon*, commonly found in Hokkaido (c).

The water samples were transported to the laboratory. While they were transported, the samples were kept at 4°C ((for up to 48 hours before filtration) to prevent eDNA degradation. The water samples were filtered and stored at -20° until eDNA extraction. DNA was extracted from the filters using a DNeasy Blood and Tissue Kit (Qiagen, Hilden, Germany). Detailed information on the DNA extraction method is available in [8] and Supplementary information.

### C. Paired-end library preparation, MiSeq sequencing and sequence data processing

Because a preliminary experiment suggested that MiMammal-U primers could not effectively amplify elephant or bear sequences, two complementary primersets for elephants and bears were included in the PCR (MiMammal-E and -B). The mixed MiMammal-U/E/B primers (hereafter, MiMammal-mix), combined with MiSeq sequencing primers and six random bases (N), was used for the multiplex PCR (see Table S4 for detailed information).

The first PCR amplified the target region using the MiMammal-mix primer set. The product of the first PCR was then purified and used as a template for the second PCR. A second PCR was performed to append MiSeq adaptors and dual index sequences to the first PCR products (Table S4). The indexed second PCR products were pooled and purified, and the DNA library was sequenced on the MiSeq platform using a MiSeq v2 Reagent Kit for 2 × 150 bp PE (Illumina, San Diego, CA, USA).

The preprocessing of the obtained sequence, OTU clustering and taxa assignments were performed using the pipeline described in Miya et al. [8]. Briefly, the overall quality of the MiSeq reads was evaluated, and the high quality pair-end reads were assembled, filtered, cleaned, and applied to the clustering process and taxonomic assignments using local BLASTN searches against a custom-made database [11]. The custom-made database was generated by downloading all mitogenome sequences from Sarcopterygii deposited in NCBI Organelle Genome Resources (http://www.ncbi.nlm.nih.gov/genomes/OrganelleResource.cgi?taxid=8287). As of 15 March 2016, the database covers 1,881 species across a wide range of families and genera. The detailed information on the PCR reaction, thermal cycle profile, purification method and sequence handling procedures is given in Supplementary information and in [8].

## III. RESULTS AND DISCUSSION

First, the performance of MiMammal-U primers was evaluated *in silico* and using 25 extracted mammalian DNA samples and eDNA extracted from water samples from zoo cages (for which the mammal species composition was exactly known). All of the results suggested that MiMammal primers were capable of detecting diverse groups of mammals from eDNA samples (Supplementary information).

As a result of MiSeq sequencing and data pre-processing, five samples from primary forest ponds generated 75,214 reads (Table 1). From all of these samples, house mouse (*Mus musculus*) sequences were frequently detected. From Pond 2 in Teshio Forest, sequences of grey-sided vole (*Myodes rufocanus*), Norway rat (*Rattus norvegicus*) and long-clawed shrew (*Sorex unguiculatus*), which are common small mammals in this forest (Table S5), were detected. From two ponds in Tomakomai forest, Sika deer (*Cervus nippon*) and common raccoon (*Procyon lotor*) sequences were detected, and these species are indeed common in this area (Table S5). In general, our method successfully detected sequences of terrestrial mammals inhabiting the sampling area.

**Table 1.** Sequence counts of detected species from water samples collected in forest ponds in Hokkaido, Japan

| Detected species (common name) | Species name | Teshio Pond 1 | Teshio Pond 2 | Teshio Pond 2 | Tomakomai Pond 1 | Tomakomai Pond 2 |
| --- | --- | --- | --- | --- | --- | --- |
| Sika deer | *Cervus nippon* | 0 | 0 | 0 | 19,452 | 1,306 |
| House mouse | *Mus musculus* | 1,171 | 128 | 701 | 127 | 846 |
| Grey red-backed vole | *Myodes rufocanus* | 0 | 497 | 393 | 0 | 0 |
| Common raccoon | *Procyon lotor* | 0 | 0 | 0 | 741 | 1,831 |
| Norway rat | *Rattus norvegicus* | 0 | 873 | 0 | 0 | 0 |
| Long-clawed shrew | *Sorex unguiculatus* | 0 | 0 | 181 | 0 | 0 |
| Other sequences |  | 6,740 | 3,419 | 156 | 16,453 | 20,199 |
| Total sequence count |  | 7,911 | 4,917 | 1,431 | 36,773 | 24,182 |
| Volume of 2nd PCR product added to the final library (μl) | | 20 | 20 | 20 | 10 | 10 |

Although this study demonstrated the usefulness of terrestrial water as a trapping medium of eDNA, several issues should be further addressed in future studies. For example, the present study as well as a previous study [7] used water samples to collect eDNA, but this might not be the most effective approach. Artificial water that attracts mammals (e.g., water with a high concentration of nutrients) might be a more effective trapping medium for mammalian eDNA. In addition, the relationship between the number of reads and the abundance of mammals in a system is less clear than that in aquatic/semiaquatic systems because the input of mammalian eDNA relies on contacts of mammals with terrestrial water. Furthermore, issues such as cross-contaminations and false positive detection should be carefully examined, as discussed in previous studies [8]) as well as Supplementary information.

## IV. CONCLUSION

In the present study, we showed that eDNA combined with our new primer sets and the MiSeq platform can be a useful tool for detecting mammals living in a terrestrial ecosystem. Describing and monitoring mammal diversity in a forest ecosystem is one of the critical steps in biodiversity conservation and management, but it is laborious and costly if one relies on traditional survey methods such as direct visual census and automated camera methods. The eDNA approach presented here is non-invasive and efficient. Moreover, it does not require setting special equipment in a field (e.g., automated cameras), and thus there is no risk of equipment loss/destruction. We propose that the eDNA metabarcoding approach could serve as an effective tool for taking snapshots of mammal biodiversity in terrestrial ecosystems.

## Ethics

This study was approved by Yokohama Zoological Gardens ZOORASIA. Water sampling in the forest ponds were obtained by Hokkaido University.

## Authors’ contributions

MU and MM conceived and designed research; MU, HF, TI, OK, KS, KM and MM performed sampling; MU, HF, TI, TS and MM performed experiments; MU, YS, MT and WI performed data analysis; MU and MM wrote the early draft and complete it with significant inputs from all authors.

## Acknowledgements

We would like to thank Mr. Noriya Saito, assistant manager of Yokohama Zoological Gardens ZOORASIA for help in sampling at the zoo, and Mr. Sho Sakurai for assistance with experiments.

## I. SUPPLEMENTARY METHODS

Please note that the water filtering, DNA extraction, PCR methods and sequence processing procedures described below were also applied to the samples collected from ponds in the primary forests.

### A. Sampling site of zoo samples

In order to test the versatility of the newly designed primers for metabarcoding eDNA from mammals, we sampled water from cages in Zoorasia Yokohama Zoological Gardens, Yokohama, Japan (35 29 42 N, 139 31 35 E; Fig.S1). We chose the zoo as a sampling site because the mammal species composition in a cage is exactly known, and because the zoo reared diverse taxonomic groups of animals (i.e., > 100 species, including many mammals and birds, were reared). Ten cages were selected as sampling places: cages of brown fur seal (*Arctocephalus pusillus*), black rhinoceros (*Diceros bicormis*), Indian elephant (*Elephas maximus*), Japanese macaque (*Macaca fuscata*), red kangaroo (*Macropus rufus*), Indian lion (*Panthera leo*), Sumatran tiger (*Panthera tigris*), Malayan tapir (*Tapirus indicus*) and polar bear (*Ursus maritimus*), and Savanna cage (in which 4 mammal species, zebra [*Equus quagga*], giraffe [*Giraffa camelopardalis*], eland [*Tragelaphus oryx*] and cheetah [*Acinonyx jubatus*], are reared in the same cage) (Fig.S1). These samples represent diverse taxonomic groups of mammals (Table S1). Water samples were collected from either a pool (for brown fur seal, Sumatran tiger and polar bear), a small bathing place (for Indian elephant and Malayan tapir), drinking water (for black rhinoceros, red kangaroo, Indian lion and Savanna cage) or a moat surrounding a cage (for Japanese macaque).

**Figure S1.**
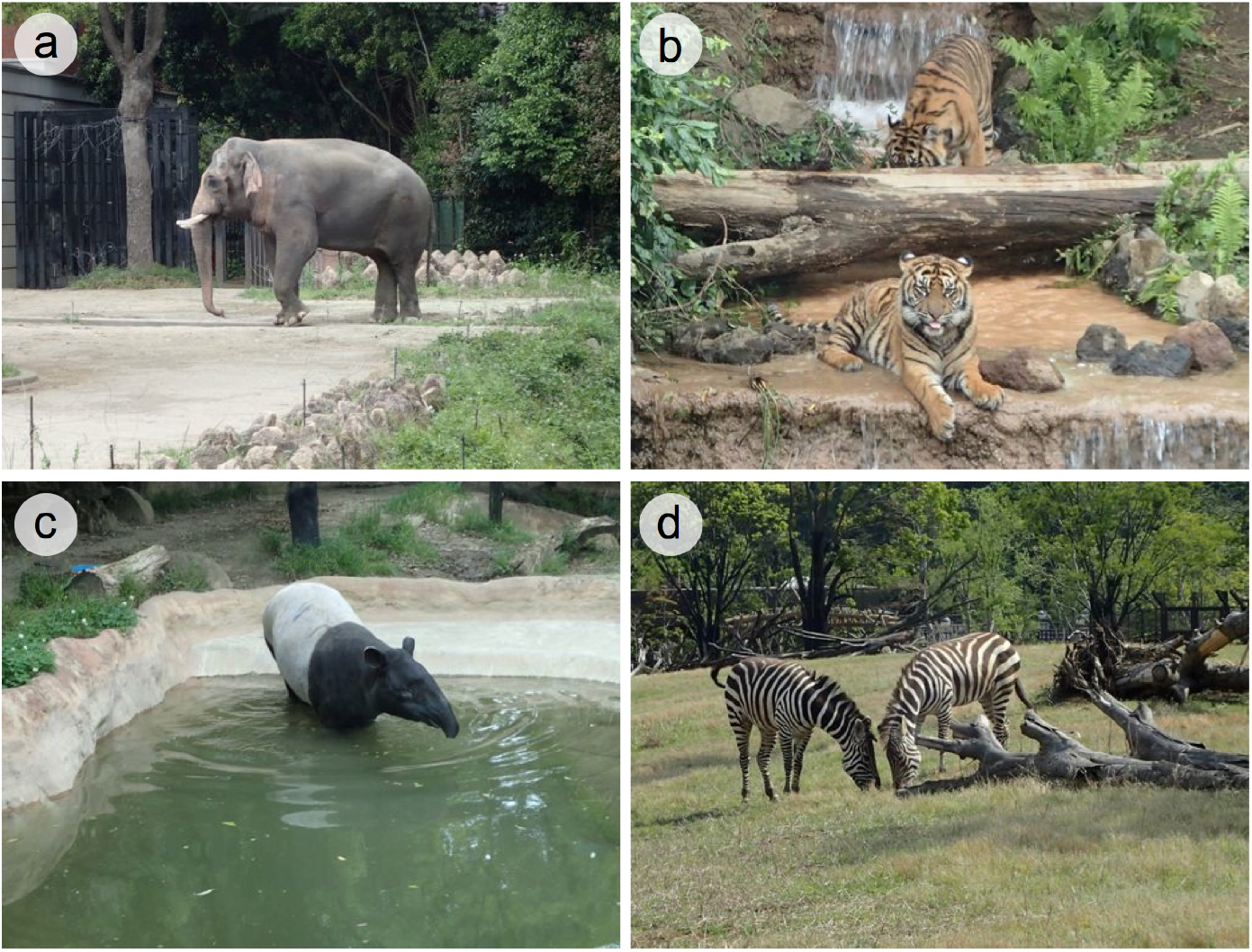
Example images of target mammal species. a-d, Mammals in the zoo cages. Indian elephant, *Elephas maximus* (a), Sumatran tiger, *Panthera tigris* (b), Malayan tapir, *Tapirus indicus* (c) and Zebra, *Equus quagga*, in the savanna cage (d)

**Table S1.** Classification of the target mammal species in the Zoorasia experiment

| Common name | Species name | Order | Familiy |
| --- | --- | --- | --- |
| Brown fur seal | *Arctocephalus pusillus* | Carnivora | Otariidae |
| Black rhinoceros | *Diceros bicornis* | Perissodactyla | Rhinocerotidae |
| Asian elephant | *Elephas maximus* | Proboscidea | Elephantidae |
| Japanese macaque | *Macaca fuscata* | Primates | Cercopithecidae |
| Red kangaroo | *Macropus rufus* | Diprotodontia | Macropodidae |
| Indian lion | *Panthera leo* | Carnivora | Felidae |
| Tiger | *Panthera tigris* | Carnivora | Felidae |
| Malayan tapir | *Tapirus indicus* | Perissodactyla | Tapiridae |
| Polar bear | *Ursus maritimus* | Carnivora | Ursidae |
| Zebra | *Equus quagga chapmani* | Perissodactyla | Equidae |
| Giraffe | *Giraffa camelopardalis* | Artiodactyla | Giraffidae |
| Eland | *Tragelaphus oryx* | Artiodactyla | Bovidae |
| Cheetah | *Acinonyx jubatus* | Carnivora | Felidae |

### B. Water sampling and DNA extraction

All sampling and filtering equipment was washed with a bleach solution before use. For water sampling in the zoo, approximately 500 ml of water was collected using a 500-ml polyethylene bottle from each sampling place (26 June 2015). The collected water samples were immediately taken back to the laboratory and filtered using 47-mm diameter glass-fibre filters (nominal pore size, 0.7 *µ*m; Whatman, Maidstone, UK). The sampling bottles were shaken vigorously before the filtration. After the filtration, each filter was wrapped in commercial aluminium foil and stored at -20°C before eDNA extraction. Five hundred ml of Milli-Q water was used as the negative control, and sampling bottles filled with Milli-Q water were treated identically to the eDNA samples in order to monitor contamination during the bottle handling, water filtering and subsequent DNA extraction.

DNA was extracted from the filters using a DNeasy Blood and Tissue Kit (Qiagen, Hilden, Germany) in combination with a spin column (EZ-10; Bio Basic, Markham, Ontario, Canada). After removing the attached membrane from the spin column (EZ-10), the filter was tightly folded into a small cylindrical shape and placed in the spin column. The spin column was centrifuged at 6,000 g for 1 min to remove redundant water from the filter. The column was then placed in a new 2-ml tube and subjected to lysis using proteinase K. Before lysis, Milli-Q water (300 *µ*l), proteinase K (10 *µ*l) and buffer AL (100 *µ*l) were mixed and the mixed solution was gently pipetted onto the folded filter in the spin column. The column was then placed on a 56°C preheated aluminium heat block and incubated for 15 min. After the incubation, the spin column was centrifuged at 6,000 g for 1 min to collect DNA. In order to increase DNA yields from the filter, 200 *µ*l of sterilized TE buffer was gently pipetted onto the folded filter and the spin column was again centrifuged at 6,000 g for 1 min. The collected DNA solution (*ca*. 500 *µ*l) was purified using a DNeasy Blood and Tissue Kit following the manufacturer's protocol.

### C. Detailed conditions for paired-end library preparation

Prior to the library preparation, work-space and equipment were sterilized, filtered pipet tips were used, and separation of pre- and post-PCR was carried out to safeguard against cross-contamination. We also employed negative controls to monitor contamination during the experiments.

The first PCR carried out with a 12-*µ*l reaction volume containing 6.0 *µ*l 2 × KAPA HiFi HotStart ReadyMix (KAPA Biosystems, Wilmington, WA, USA), 0.7 *µ*l of each primer (5 *µ*M), 2.6 *µ*l sterilized distilled H_2_O and 2.0 *µ*l template. When the first PCR was multiplexed (i.e., when it was treated using MiMammal-mix), the final concentration of each primer (MiMammal-U/E/B) was 0.1 *µ*M (0.3 *µ*M total concentration of primers). The thermal cycle profile after an initial 3 min denaturation at 95°C was as follows (35 cycles): denaturation at 98°C for 20 s; annealing at 65°C for 15 s; and extension at 72°C for 15 s, with the final extension at the same temperature for 5 min. We performed triplicate first-PCR, and the replicates were pooled in order to mitigate the PCR dropouts. The pooled first PCR products were purified using Exo-SAPIT (Affymetrix, Santa Clara, CA, USA). The pooled, purified, and 10-fold diluted first PCR products were used as templates for the second PCR.

The second-round PCR (second PCR) was carried out with a 24-*µ*l reaction volume containing 12 *µ*l of 2 × KAPA HiFi HotStart ReadyMix, 1.4 *µ*l each primer (5 *µ*M), 7.2 *µ*l sterilized distilled H_2_O and 2.0 *µ*l template. Different combinations of forward and reverse indices were used for different templates (samples) for massively parallel sequencing with MiSeq. The thermal cycle profile after an initial 3 min denaturation at 95°C was as follows (12 cycles): denaturation at 98°C for 20 s; annealing and extension combined at 65°C (shuttle PCR) for 15 s with the final extension at 72°C for 5 min. The products of the second PCR were combined, purified, excised and sequenced on the MiSeq platform using a MiSeq v2 Reagent Kit for 2 × 150 bp PE.

**Table S2.**
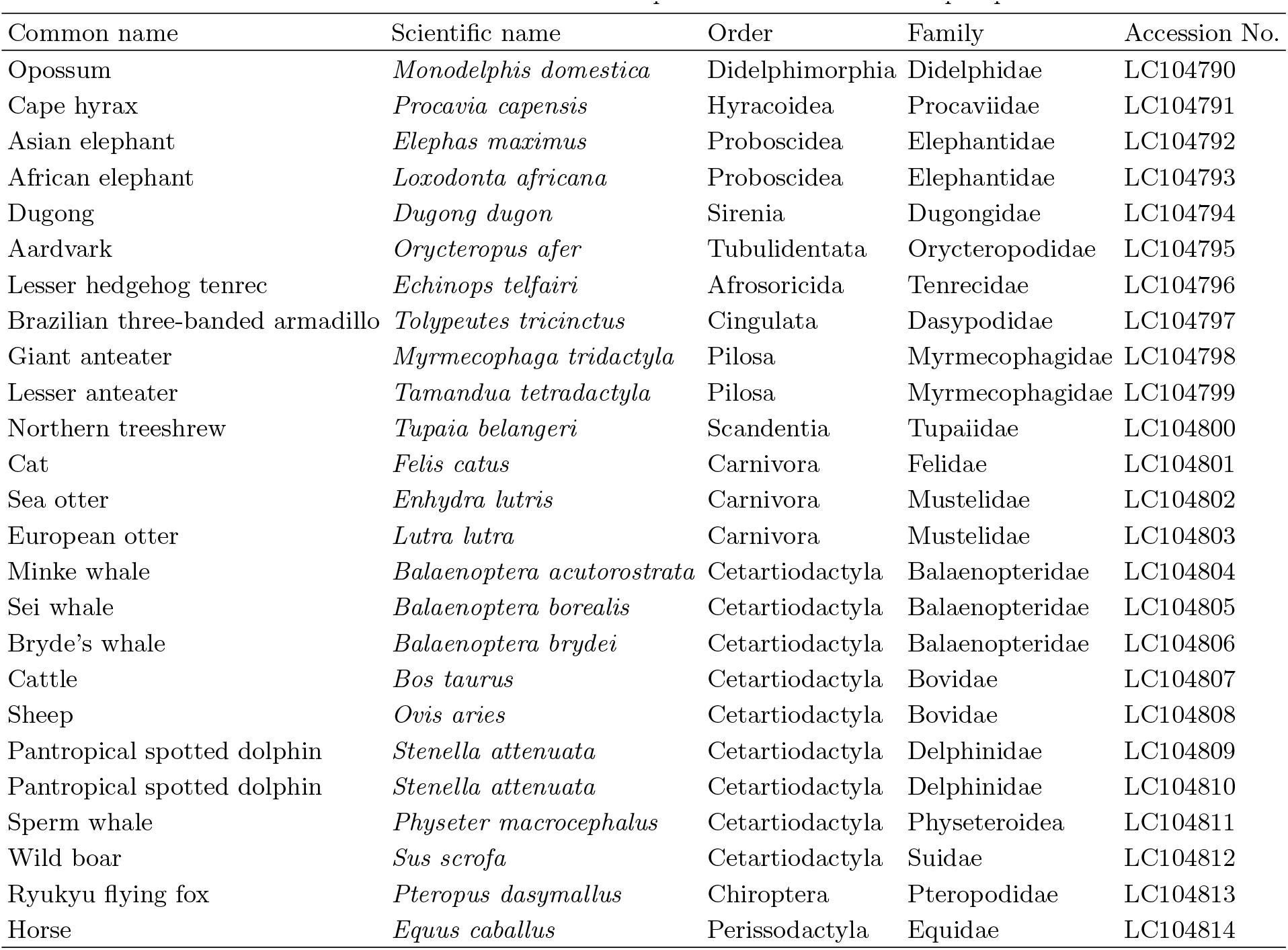
Extract DNAs used to test the performance of the developed primer set

### D. Sequence read processing and taxonomic assignment

The overall quality of the MiSeq reads was evaluated, and the reads were assembled using the software FLASH with a minimum overlap of 10 bp [1]. The assembled reads were further filtered and cleaned, and the pre-processed reads were subjected to the clustering process and taxonomic assignments. The pre-processed reads from the above custom pipeline were dereplicated using UCLUST [2]. Those sequences represented by more at least 10 identical reads were subjected to the downstream analyses, and the remaining under-represented sequences (with less than 10 identical reads) were subjected to pairwise alignment using UCLUST. If the latter sequences (observed from less than 10 reads) showed at least 99% identity with one of the former reads (i.e., no more than one or two nucleotide differences), they were operationally considered as identical (owing to sequencing or PCR errors and/or actual nucleotide variations in the populations).

The processed reads were subjected to local BLASTN searches against a custom-made database [3]. The custom-made database was generated as described in the main text. The top BLAST hit with a sequence identity of at least 97% and E-value threshold of 10^-5^ was applied to species assignments of each representative sequence. The detailed information for the above bioinformatics pipeline from data pre-processing through taxonomic assignment is available in the supplemental information in [4].

## II. SUPPLEMENTARY RESULTS

### A. Tests of versatility of new primers with DNA extracted from tissue and *in silico*

The performance of MiMammal-U primers was evaluated using 25 extracted mammalian DNA samples representing major groups of mammals (Table S2). All DNAs were successfully amplified, and the sequences were deposited as shown in Table S2. In addition, levels of interspecific variation in the target region was computationally evaluated using downloaded sequences of 741 mammal species, and the results of this evaluation suggested that these primers designed here were capable of distinguishing among mammals in diverse groups of mammals.

**Table S3.**
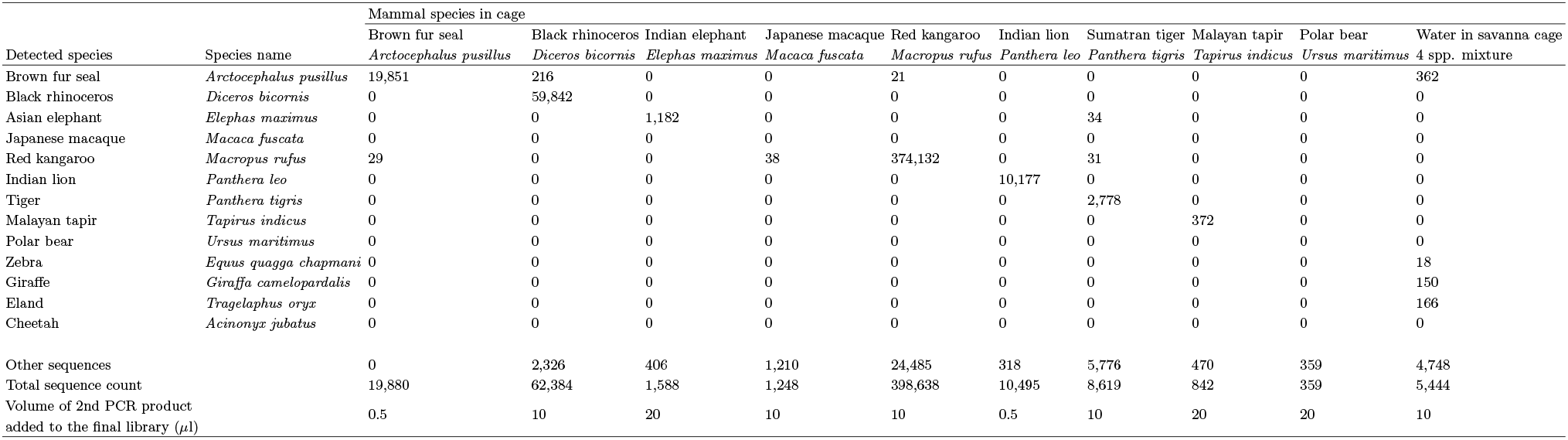
Sequence reads of detected species from water samples collected in the zoo

| Detected species | Species name | Mammal species in cage |  |  |  |  |  |  |  |  |  |
| --- | --- | --- | --- | --- | --- | --- | --- | --- | --- | --- | --- |
|  |  | Brown fur seal | Black rhinoceros | Indian elephant | Japanese macaque | Red kangaroo | Indian lion | Sumatran tiger | Malayan tapir | Polar bear | Water in savanna cage |
|  |  | *Arctocephalus pusillus* | *Diceros bicornis* | *Elephas maximus* | *Macaca fuscata* | *Macropus rufus* | *Panthera leo* | *Panthera tigris* | *Tapirus indicus* | *Ursus maritimus* | 4 spp. mixture |
| Brown fur seal | *Arctocephalus pusillus* | 19,851 | 216 | 0 | 0 | 21 | 0 | 0 | 0 | 0 | 362 |
| Black rhinoceros | *Diceros bicornis* | 0 | 59,842 | 0 | 0 | 0 | 0 | 0 | 0 | 0 | 0 |
| Asian elephant | *Elephas maximus* | 0 | 0 | 1,182 | 0 | 0 | 0 | 34 | 0 | 0 | 0 |
| Japanese macaque | *Macaca fuscata* | 0 | 0 | 0 | 0 | 0 | 0 | 0 | 0 | 0 | 0 |
| Red kangaroo | *Macropus rufus* | 29 | 0 | 0 | 38 | 374,132 | 0 | 31 | 0 | 0 | 0 |
| Indian lion | *Panthera leo* | 0 | 0 | 0 | 0 | 0 | 10,177 | 0 | 0 | 0 | 0 |
| Tiger | *Panthera tigris* | 0 | 0 | 0 | 0 | 0 | 0 | 2,778 | 0 | 0 | 0 |
| Malayan tapir | *Tapirus indicus* | 0 | 0 | 0 | 0 | 0 | 0 | 0 | 372 | 0 | 0 |
| Polar bear | *Ursus maritimus* | 0 | 0 | 0 | 0 | 0 | 0 | 0 | 0 | 0 | 0 |
| Zebra | *Equus quagga chapmani* | 0 | 0 | 0 | 0 | 0 | 0 | 0 | 0 | 0 | 18 |
| Giraffe | *Giraffa camelopardalis* | 0 | 0 | 0 | 0 | 0 | 0 | 0 | 0 | 0 | 150 |
| Eland | *Tragelaphus oryx* | 0 | 0 | 0 | 0 | 0 | 0 | 0 | 0 | 0 | 166 |
| Cheetah | *Acinonyx jubatus* | 0 | 0 | 0 | 0 | 0 | 0 | 0 | 0 | 0 | 0 |
| Other sequences |  | 0 | 2,326 | 406 | 1,210 | 24,485 | 318 | 5,776 | 470 | 359 | 4,748 |
| Total sequence count |  | 19,880 | 62,384 | 1,588 | 1,248 | 398,638 | 10,495 | 8,619 | 842 | 359 | 5,444 |
| Volume of 2nd PCR product added to the final library (μl) | | 0.5 | 10 | 20 | 10 | 10 | 0.5 | 10 | 20 | 20 | 10 |

**Table S4.**
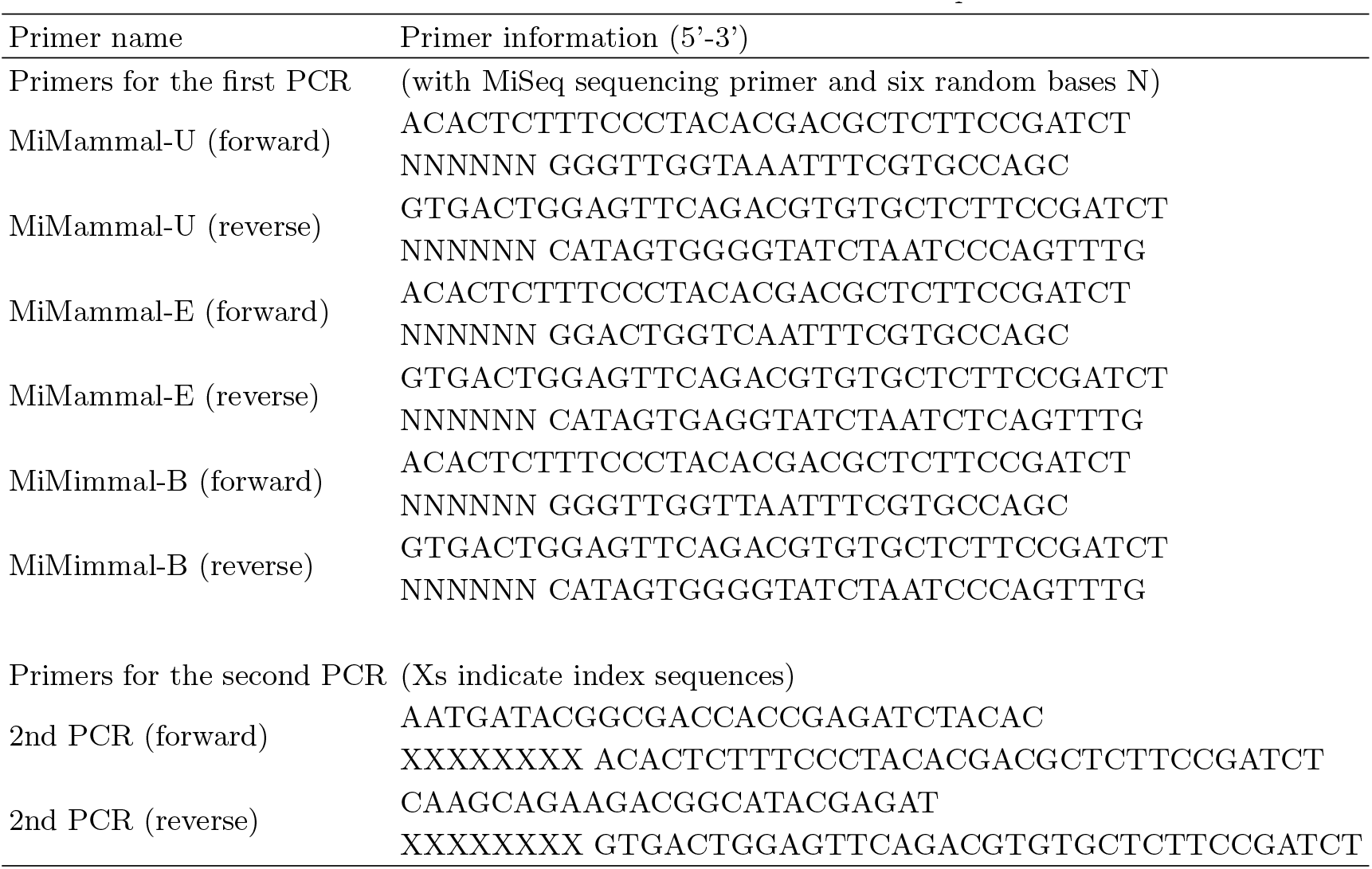
Detailed information for MiMammal primers

### B. Detection of target species from zoo cages

As a result of MiSeq sequencing and data processing, 10 samples generated 509,497 sequences (Table S3). Among the 10 cages (13 species in total) examined, 10 species were successfully detected. For brown fur seal (*Arctocephalus pusillus*), black rhinoceros (*Diceros bicornis*), Indian elephant (*Elephas maximus*), red kangaroo (*Macrofus rufus*), Indian lion (*Panthera leo*), Sumatran tiger (*Panthela tigris*) and Malayan tapir (*Tapirus indicus*), large proportions of the total sequence counts were the target species origin (at least > 30%, Fig. S2). These mammals frequently contacted the waters in the cages, and such behaviours are likely to influence the numbers (and proportions) of target species sequences detected in the samples. For the Savanna cage sample, only 6.6% of total sequences were of target species (zebra, giraffe, and eland) origin. The Savanna cage occupies a relatively large area, and the mammals in the cage do not frequently contact the water. The large cage area and the animals’ behaviour might have resulted in the low proportion of target species sequences from this cage.

In contrast to the above-mentioned target species, we could not detect the sequences of cheetah (*Acinonyx jubatus*), Japanese macaque (*Macaca fuscata*) and polar bear (*Ursus maritimus*). For cheetah, the reason for the non-detection seems to lie in the animal's behaviour. The cheetah is often resting on a tree far from the water, and it has much less opportunity to contact the water compared to the other mammals (zebra, giraffe and eland) in the cage. For Japanese macaque, the water sample was taken from a moat surrounding the cage. This means that the Japanese macaques living in the cage cannot directly contact the water. Therefore, non-detection of the Japanese macaque sequences from the water sample may not be surprising. Conversely, direct contacts with the water might dramatically increase the chance of the detection of mammals with the eDNA approach. For polar bear, the pool in the cage is always sterilized with a low concentration of chloride (< 0.05%) in order to prevent harmful algal blooms, and this chloride might degrade DNA from the polar bear. In addition, the pool was intensively washed after completely removing the water from the pool one day before the water sampling. The addition of chloride and the intensive washing was likely to have degraded/removed the polar bear eDNA, which probably prevented the detection here of polar bear sequences.

**Figure S2.**
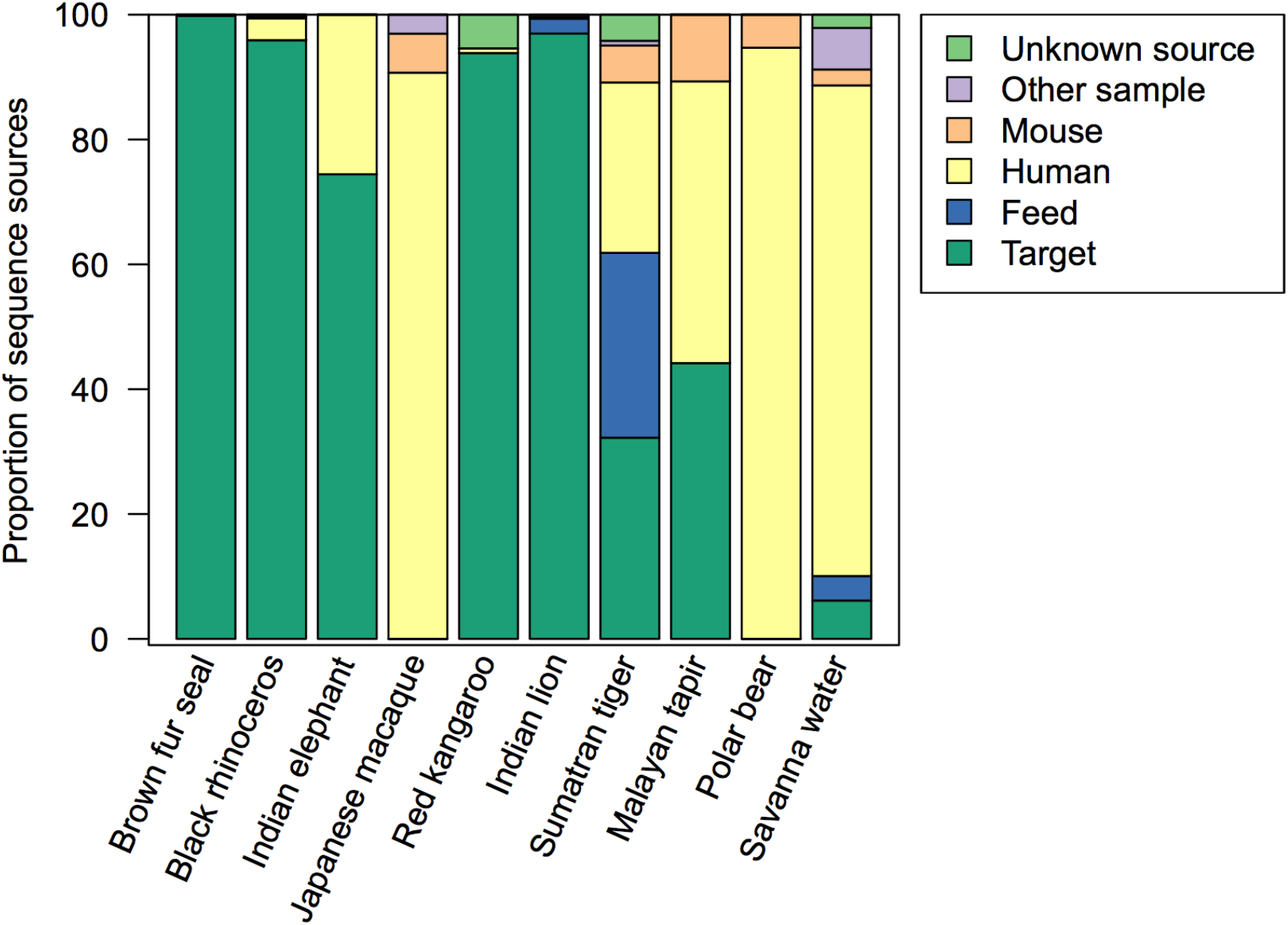
Proportions of sequence sources of the zoo samples. Different colours indicate different sources as described in the legend.

Another possible cause of the non-detection of some mammal species may have lain in the experimental conditions (e.g., PCR conditions). To examine this possibility, we modified the annealing temperature and primer concentrations of the first PCR. After several trial-and-error attempts, we detected 1,400 reads of Japanese macaque sequence (from 13,259 total reads) from the Japanese macaque water sample and 187 reads of cheetah sequence (from 51,118 total reads) from the savanna water sample under the conditions of 62°C annealing temperature and a total 15 *µ*M MiMammal-mix primer set (each primer at 5 *µ*M). We also detected 6,990 reads of polar bear sequence (from 61,347 total reads) from the polar bear pool sample under the conditions of annealing temperature 62°C and 5 *µ*M MiMammal-B primer set. Altogether, MiMammal primers successfully detected all mammals examined in the zoo cages, but the detection depended on the experimental conditions.

**Table S5.** List of common mammals in Teshio and Tomakomai Experimental Forests

| Common name | Scientific name | Detection by eDNA |
| --- | --- | --- |
| **Teshio Experimental Forest** |  |  |
| Eurasian common shrew | *Sorex caecutiens* Laxmann, 1788 |  |
| Long-clawed shrew | *Sorex unguiculatus* Dobson, 1890 | detected |
| Gray red-backed vole | *Myodes rufocanus* Sundevall, 1846 | detected |
| Large Japanese field mouse | *Apodemus speciosus* Temminck, 1844 |  |
| Small Japanese field mouse | *Apodemus argenteus* Temminck, 1844 |  |
| Norway rat | *Rattus norvegicus* Berkenhout, 1769 | detected |
| Siberian chipmunk | *Tamias sibiricus* Laxmann, 1769 |  |
| Red fox | *Vulpes vulpes* Linnaeus, 1758 |  |
| Raccoon dog | *Nyctereutes procyonoides* Gray, 1834 |  |
| Raccoon | *Procyon lotor* Linnaeus, 1758 |  |
| Brown bear | *Ursus arctos* Linnaeus, 1758 |  |
| Sika deer | *Cervus Nippon* Temminck, 1836 |  |
| **Tomakomai Experimental Forest** |  |  |
| Eurasian common shrew | *Sorex caecutiens* Laxmann, 1788 |  |
| Long-clawed shrew | *Sorex unguiculatus* Dobson, 1890 |  |
| Gray red-backed vole | *Myodes rufocanus* Sundevall, 1846 |  |
| Large Japanese field mouse | *Apodemus speciosus* Temminck, 1844 |  |
| Small Japanese field mouse | *Apodemus argenteus* Temminck, 1844 |  |
| Siberian chipmunk | *Tamias sibiricus* Laxmann, 1769 |  |
| Raccoon dog | *Nyctereutes procyonoides* Gray, 1834 |  |
| Raccoon | *Procyon lotor* Linnaeus, 1758 | detected |
| Sika deer | *Cervus Nippon* Temminck, 1836 | detected |

### C. Detection of non-target species from zoo cages

In the zoo experiment, the most frequently detected non-target species were human and mouse (Fig. S2). These species are ubiquitous in the sampling area, and they are likely to have frequent opportunities to directly/indirectly contact the water. Thus, it is difficult to distinguish the origin of the sequences, i.e., whether they came from the sampling site and/or contaminations during the experimental procedures. In addition, feeds of the target species were often detected (Fig. S2), which indicates that eDNA from dead organisms can also be easily detected.

As already pointed out in a previous study [4], the most serious pitfall of eDNA is the risk of contamination. To avoid this risk, we performed standard decontamination procedures of the laboratory spaces and equipment (e.g., routine cleaning of experiment benches using DNA remover, the use of filter-tips, and physical separation of DNA extraction and PCR rooms). Despite these efforts, on average, approximately 10% of the total number of reads was considered to be originate from contamination (i.e., sequences were from unknown origin and/or other samples, Fig. S2). This rate of contamination is in a similar range as that of other studies using the metabarcoding technique. Such contamination issues are among the most challenging problems that still remained unsolved regarding experimental applications of the metabarcoding technique.

### D. List of common mammals caught by box trap or captured by automated cameras

The mammalian fauna had been monitored in Teshio and Tomakomai Experimental Forests of Hokkaido University using box traps and automated cameras from 2009 to 2015. The results shown in Table S5 indicate that the mammals detected by the eDNA metabarcoding approach are indeed common in the present water-sampling area in Hokkaido, Japan.

